# Absence of posterior commissure and sub-commissural organ precedes encephalocele development in a new mouse model

**DOI:** 10.1101/2025.11.10.687701

**Authors:** Hiu Nam Chan, Dawn Savery, Shreeta Chakraborty, Nina Wenzlitschke, Pedro P. Rocha, Andrew J. Copp

## Abstract

Encephalocele is a congenital defect involving herniation of the meninges, with or without brain tissue, outside the skull. Although traditionally considered a neural tube defect (NTD) alongside anencephaly and open spina bifida (myelomeningocele), encephalocele typically shows well-formed brain tissue, and is likely to represent a later-arising, post-neurulation developmental anomaly, with a different pathogenic mechanism from the ‘open’ NTDs. A detailed understanding of encephalocele pathogenesis requires experimental studies and, recently, we developed a new mouse model in which the genes encoding FGF3, 4 and 15 are over-expressed in the embryonic day (E) 9.5 brain. Encephalocele subsequently develops as a broad forebrain-midbrain swelling, visible from E11.5, that resolves by birth into a focal brain herniation resembling human parieto-occipital encephalocele. A structural analysis of the brain in mutant embryos reveals absence of the posterior commissure and sub-commissural organ, and at later stages the pineal gland. These structures normally develop just rostral to the forebrain-midbrain boundary, and Pax6 immuno-histochemistry demonstrates that this boundary remains intact in the mutant embryos. Histological analysis reveals a more general change in tissue composition of the neural tube roof in the forebrain-midbrain region, with diminished thickness of the neuroepithelium and increased thickness of the overlying layer, including the non-neural ectoderm (future epidermis). We conclude that the posterior commissure and sub-commissural organ, previously implicated in hydrocephalus, may also be fundamental for development of the earlier-arising brain malformation, encephalocele.

## INTRODUCTION

Neural tube defects (NTDs) are congenital nervous system malformations that arise during embryonic development due to anomalous development of the brain and/or spinal cord. Failed closure of the neural tube leads to ‘open’ NTDs (e.g. anencephaly, myelomeningocele) whereas ‘closed’ NTDs are conditions in which the neural tube completes closure but subsequently becomes abnormal, with skin covering [1, 2].

Encephalocele is a closed NTD in which the meninges herniate outside the skull, with or without enclosed brain tissue. This condition can affect fronto-ethmoidal, parieto-occipital and peri-torcular locations along the skull midline [3]. The median prevalence of encephalocele is 1-3 per 10,000 births (i.e. approximately 10% of all NTDs), with considerable variation between human populations [4]. Although usually sporadic, encephalocele can be syndromic, as in Meckel syndrome, an autosomal recessive ciliopathy that commonly includes occipital encephalocele [5].

Individuals with encephalocele usually undergo surgery in the newborn period, with removal of the herniated meninges and neural tissue, water-tight closure of the dura and repair of the skull defect. An individual’s subsequent prognosis depends on several factors including size, location and the herniated content of the encephalocele, with neurological problems such as seizures and ataxia being common in children post-surgery [6].

The embryonic events that precede encephalocele are largely unknown, in contrast to the pathogenic mechanisms of open NTDs, which have been extensively studied in animal models [7]. Encephaloceles have been variously reported as primary neurulation defects [8], or defects of skull development with secondary brain herniation [9]. Several mouse models of frontal encephalocele have been described: *forebrain overgrowth* (*fog*) which results from loss of *Apaf1* gene function, with mis-regulation of programmed cell death [10], and *tuft*, a mutation of the *Tet1* gene, that encodes an epigenetic regulator [11]. Both have been attributed to faulty neural tube closure in the forebrain region. In contrast, mice homozygous for the *Grhl2^Axd^* mutation develop either frontal encephalocele or brain protrusions through the skull base [12], with no evidence to implicate faulty neural tube closure in either case. It would appear that the embryonic pathogenesis of frontal encephalocele is deserving of further detailed study to clarify the underlying mechanisms.

Models of the commonest human encephalocele variant that affects the parieto-occipital region, are scarce. Recently, an opportunity to investigate the pathogenesis of this encephalocele type arose following the finding of parieto-occipital encephalocele in a new mouse genetic model [13]. This model targets CCCTC-binding factor (CTCF) sites upstream of the *FGF 3, 4* and *15* genes on mouse chromosome 7. Heterozygous deletion of four CTCF binding sites (C1-C4) produced encephalocele, and this deletion was subsequently rendered conditional using the Cre/lox system [13]. CTCF is a conserved 11-zinc finger protein that functions in creating boundaries between chromatin domains. Deletion of sites C1-4 leaves the *FGF* genes uninsulated from upstream enhancers and their resulting increased expression in the embryonic day (E) 9.5 midbrain and forebrain reproduces the expression pattern of *Ano1*, a gene immediately upstream of the C1-C4 sites [13].

Here, we hypothesised that over-activation of *FGF* gene expression in this mouse model leads to an encephalocele phenotype via earlier disruption of brain development. The aim of our study was to examine the structure of the brain in a staged series of embryos and fetuses with recombination of the floxed *C1-C4* mutant construct in the dorsal brain midline. Recombination was achieved using either *Wnt1^Cre^* or *Pax3^Cre^*, both expressed in the dorsal midline, which each yield a closely similar brain herniation phenotype. Strikingly, this approach reveals a complete absence of key posterior forebrain structures: the posterior commissure, sub-commissural organ and pineal gland in the pre-encephalocele embryos, despite maintenance of the forebrain-midbrain boundary, as indicated by Pax6 expression. Moreover, a more general thinning of the dorsal midline neuroepithelium, and abnormal thickening of the overlying future skin/skeletal layer, were observed in mutant embryos and fetuses. These findings provide a starting point for an in-depth analysis of encephalocele pathogenesis in one of the first mouse models of this condition to be described.

## RESULTS

Embryos and fetuses were collected from matings between *Wnt1^Cre/+^* males and *C1-C4^flx/flx^* females, at E9.5 to E19.5, to investigate the onset of phenotypic differences between *Wnt1^+/+^; C1-C4^flx/+^* (hereafter: WT) and *Wnt1^Cre/+^; C1-C4^flx/+^* (hereafter: Het). The relative frequencies of WT and Het individuals in these litters conformed closely to the 1:1 Mendelian expectation (Table 1).

**Table 1.** Numbers of embryos and fetuses by genotype at different developmental stages from matings of *Wnt1^Cre/+^* or *Pax3^Cre/+^* males with *C1-C4^flx/flx^* females.

| Mating type | Stage | No. litters | No. embryos and fetuses |  |  |
| --- | --- | --- | --- | --- | --- |
|  |  |  | Total | WT * | Het ** |
| *Wnt1*<sup>Cre/+</sup> x | E9.5 | 1 | 10 | 4 | 6 |
| *C1-C4*<sup>flx/flx</sup> | E10.5 | 1 | 10 | 6 | 4 |
|  | E11.5 | 1 | 10 | 4 | 6 |
|  | E12.5 | 6 | 72 | 39 | 33 |
|  | E13.5 | 1 | 9 | 4 | 5 |
|  | E14.5 | 3 | 28 | 15 | 13 |
|  | E15.5 | 1 | 9 | 3 | 6 |
|  | E16.5 | 3 | 27 | 11 | 16 |
|  | E17.5 | 1 | 8 | 6 | 2 |
|  | E18.5 | 1 | 10 | 7 | 3 |
|  | E19.5 | 1 | 10 | 8 | 2 |
|  | **Totals** | **20** | **203** | **107** | **96** |
| *Pax3*<sup>Cre/+</sup> x |  |  |  |  |  |
| *C1-C4*<sup>flx/flx</sup> | E12.5 | 2 | 26 | 12 | 14 |

\* Cre-negative, *C1-C4*<sup>flx/+</sup> embryos and fetuses
\*\* Cre-positive, *C1-C4*<sup>flx/+</sup> embryos and fetuses

### Time-course of encephalocele development in *Wnt1^Cre/+^; C1-C4^flx/+^* mice

Observations at E9.5 and E10.5 (Figure 1A-D) confirmed that neural tube closure is complete in the cranial region of Het embryos (Figure 1A-F). The encephalocele phenotype becomes clearly evident from E11.5, when the posterior forebrain and midbrain region bulge superiorly (Figure 1E, F), although some Het embryos already show slight brain expansion at E10.5 (Figure 1C, D). From E11.5 to E14.5, the brain continues to expand superiorly and posteriorly, with a translucent appearance suggesting a fluid-filled space (Figure 1E-L). However, from E15.5 onwards, the expanded region narrows rostro-caudally, becoming a focal head extension that is no longer translucent in appearance. Indeed, extruded brain tissue can be seen within the herniated region (Figure 1M-V). At E19.5, just before birth, internal bleeding is seen within the encephalocele (Figure 1U, V). In some cases, the herniated sac burst during removal of late-stage fetuses from the uterus, leaving a hole in the top of the head, which corresponds to the incomplete skull vault formation, as described in Het fetuses previously [13].

**Figure 1.**
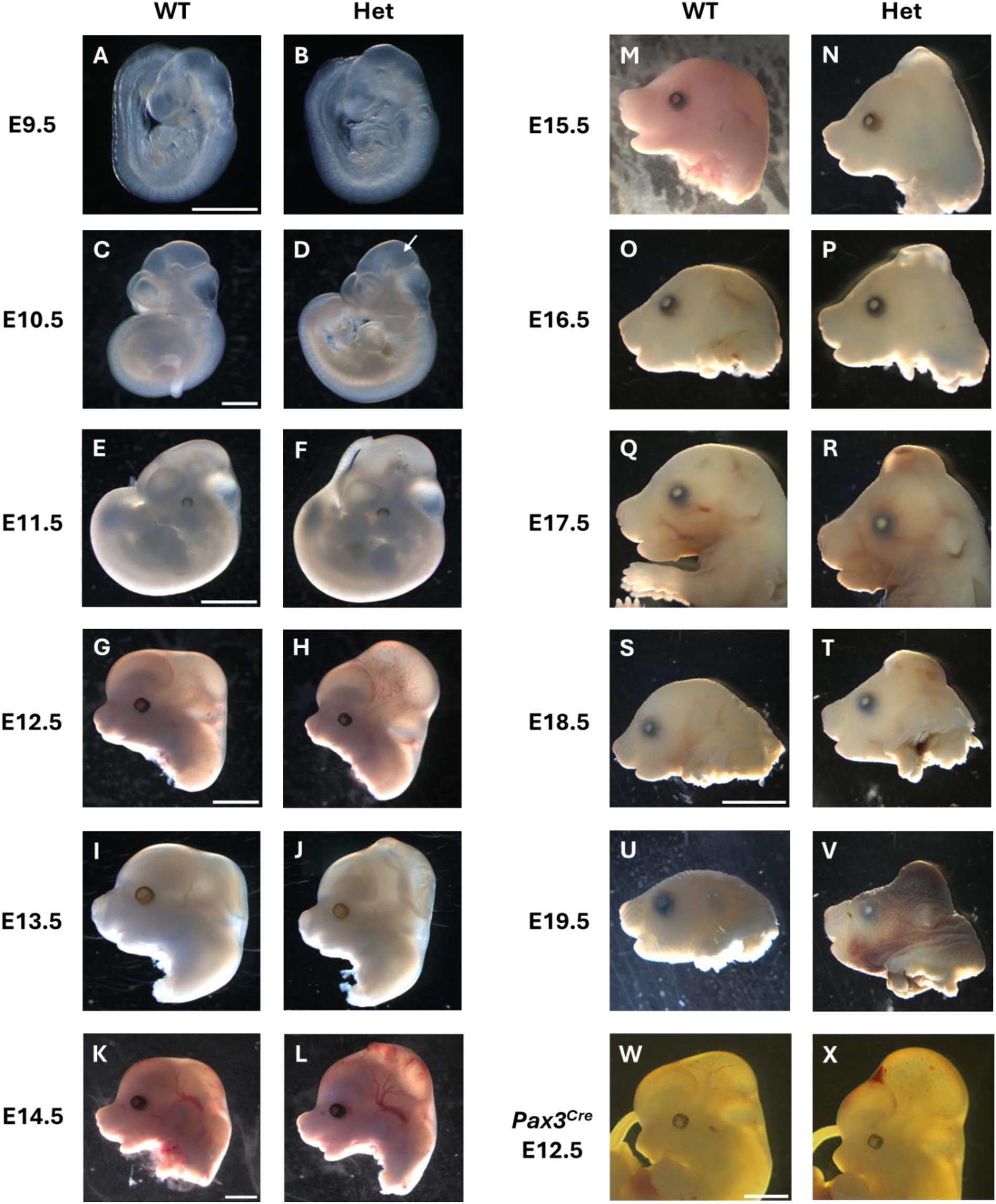
Emergence of the encephalocele phenotype across the range of embryonic and fetal stages. Left-sided views of whole embryos (A-F) or embryonic and fetal heads (G-X), comparing *Wnt1^+/+^; C1-C4^flx/+^* (WT) and *Wnt1^Cre^; C1-C4^flx/+^* (Het) individuals from E10.5 to E19.5 (A-V). E12.5 *Pax3^+/+^; C1-C4^flx/+^* (W; WT) and *Pax3^Cre/+^; C1-C4^flx/+^* (X; Het) embryos are also shown. Swelling of the posterior forebrain and midbrain first appears at E10.5 or E11.5 (arrow in D) and continues as a dorsal, translucent brain enlargement from E11.5 to E14.5 (E-L; W,X). From E15.5 (M, N), the encephalocele takes on a more focal, opaque morphology, with visible protrusion of brain tissue in the herniated region. Scale bars: 1 mm (A,B) (C-D), 2 mm (E,F) (G-J) (K-R) (W,X), 5 mm (S-V). Sample size: n = 3 or more per stage and genotype, except n = 2 for Hets at E17.5 and E19.5.

A small number of E12.5 embryos were also obtained from *Pax3^Cre/+^* x *C1-C4^flx/flx^* matings, with the Cre-positive embryos exhibiting a pre-encephalocele phenotype (Figure 1W,X), closely similar that seen in *Wnt1^Cre/+^* x *C1-C4^flx/+^* embryos. Hence, the encephalocele phenotype is not specific to a Cre driver, but rather results from recombination of the floxed *C1-C4* sites in the dorsal brain midline.

### Histological comparison of WT and Het brain development

To examine the internal morphology of encephalocele development, the brains of E10.5 to E16.5 WT and Het embryos and fetuses, from *Wnt1^Cre/+^* x *C1-C4^flx/+^* matings, were sectioned in the mid-sagittal plane and stained with H&E. At E10.5, no structural differences between WT and Het could be identified (Figure 2A-D). At E11.5, the posterior commissure (PC) first becomes visible in WT embryos as a row of cross-sectioned fibre bundles, near the basal edge of the neuroepithelium (Figure 2E,F). The PC is the first commissure to appear during brain development [14] and comprises axonal bundles that cross the midline at the level of the diencephalon, connecting equivalent parts of the two cerebral hemispheres [15]. By E12.5, the PC becomes more obvious in WT embryos, and the sub-commissural organ (SCO), a region of columnar cells immediately beneath the PC, is also detectable (Figure 2I,J). From E13.5 onwards, an additional structure, the pineal gland (PG), is visible in WT sections (Figure 2M,N). At these stages, the PG comprises a caudally-directed out-pocketing of the forebrain neuroepithelium, immediately rostral to the PC. At E14.5 and E16.5, the PC-SCO-PG forms a prominent complex in the roof of the diencephalon of WT fetuses (Figure 2Q,R,U,V).

**Figure 2.**
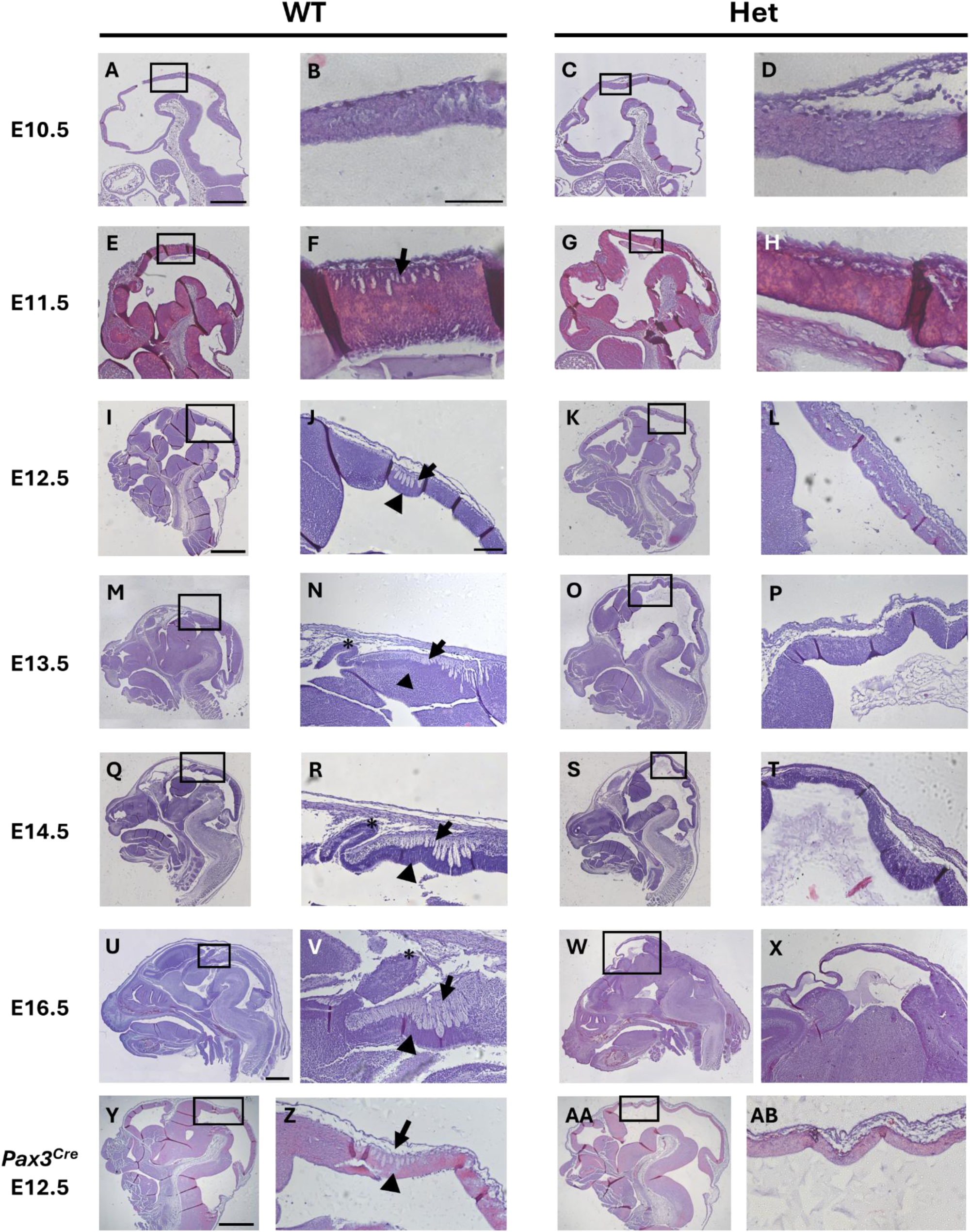
Absence of PC-SCO-PG complex in the brains of *Wnt1^Cre^; C1-C4^flx/+^* and *Pax3^Cre^; C1-C4^flx/+^* embryos. **(A-X)** H&E staining of mid-sagittal head sections of stage-matched *Wnt1^+/+^; C1-C4^flx/+^* (WT; A,E,I,M,Q,U) and *Wnt1^Cre/+^; C1-C4^flx/+^* (Het; B) embryos and fetuses, with magnified views of boxed regions at the forebrain-midbrain boundary. *Pax3^+/+^; C1-C4^flx/+^* (Y,Z; WT) and *Pax3^Cre/+^; C1-C4^flx/+^* (AA,AB; Het) embryos are also shown. Anterior is to the left. The dorsal neuroepithelium appears undifferentiated in both WT and Het at E10.5 (A-D) whereas, in WT brains from E11.5, the posterior commissure (PC) becomes visible and progressively enlarges, appearing as a row of cross-sectioned fibre tracts adjacent to the basal neuroepithelial surface (arrows in E,F,I,J,M,N,Q,R,U,V,Y,Z). Also visible in WT brains are the sub-commissural organ (SCO; arrowheads), located beneath the PC from E12.5, and the pineal gland (asterisks) as an outpouching of the neuroepithelium, from E13.5 (M,N,Q,R,U,V). In contrast, Het brains show no sign of PC, SCO or pineal gland at any stage from E11.5-E16.5 (G,H,K,L,O,P,S,T,W,X,AA,AB), with the corresponding neuroepithelium appearing undifferentiated. Scale bars: 0.5 mm (A,C,E,G), 1 mm (I,K,M,O,Q,S)(U,W)(Y,AA), 0.2 mm (B,D,F,H)(J,L,N,P,R,T,V,X,Z,AB). Sample size: n = 3 (E12.5/14.5/16.5), n = 1 (E10.5/11.5/13.5) per stage and genotype.

In striking contrast to our observations on WT brain sections, we did not observe a PC, SCO or PG in the Het caudal forebrain at any point in the E11.5-E16.5 age range. Instead, the Het neuroepithelium shows no features of local differentiation, and appears persistently thin compared with WT (Figure 2G,H,K,L,O,P,D,S,T,W,X). Moreover, it tends to be thrown into folds, within the ventricular cavity, to a greater extent than the WT neuroepithelium (e.g. compare Figure 2Q and S). Similar observations were made using *Pax3^Cre/+^* x *C1-C4^flx/flx^* litters at E12.5, with WT embryos exhibiting the PC and SCO, whereas these structures were absent from Het embryos (Figure 2Y-AB).

### Forebrain-midbrain boundary is preserved in pre-encephalocele development

The PC-SCO-PG complex develops in the caudal part of the future diencephalon, just rostral to the forebrain-midbrain boundary. This boundary is marked by Pax6 which is expressed in the forebrain but not midbrain [16]. Pax6 is also required for development of the PC-SCO-PG complex, which is absent from *small eye* mutant mice that lack Pax6 [17]. We hypothesised that abnormality of Pax6 expression, and/or a shift in the forebrain-midbrain boundary could be responsible for the absence of PC-SCO-PG in the pre-encephalocele Het mice. Immunostaining of mid-sagittal brain sections for *Pax6* at E12.5 and 14.5 revealed a similar intensity of Pax6 expression in both WT and Het neuroepithelia (Figure 3A-D). Moreover, a clear boundary of Pax6 expression can be seen at a closely similar location in both genotypes. The PC-SCO-PG complex is located just within Pax6-positive (forebrain) domain in WT, whereas the neuroepithelial regions either side of the Pax6 expression boundary in Het embryos show no morphological differentiation. We conclude that the encephalocele phenotype in Het embryos is not preceded by any obvious abnormality in Pax6 expression, and that the forebrain-midbrain boundary appears to be specified correctly.

**Figure 3.**
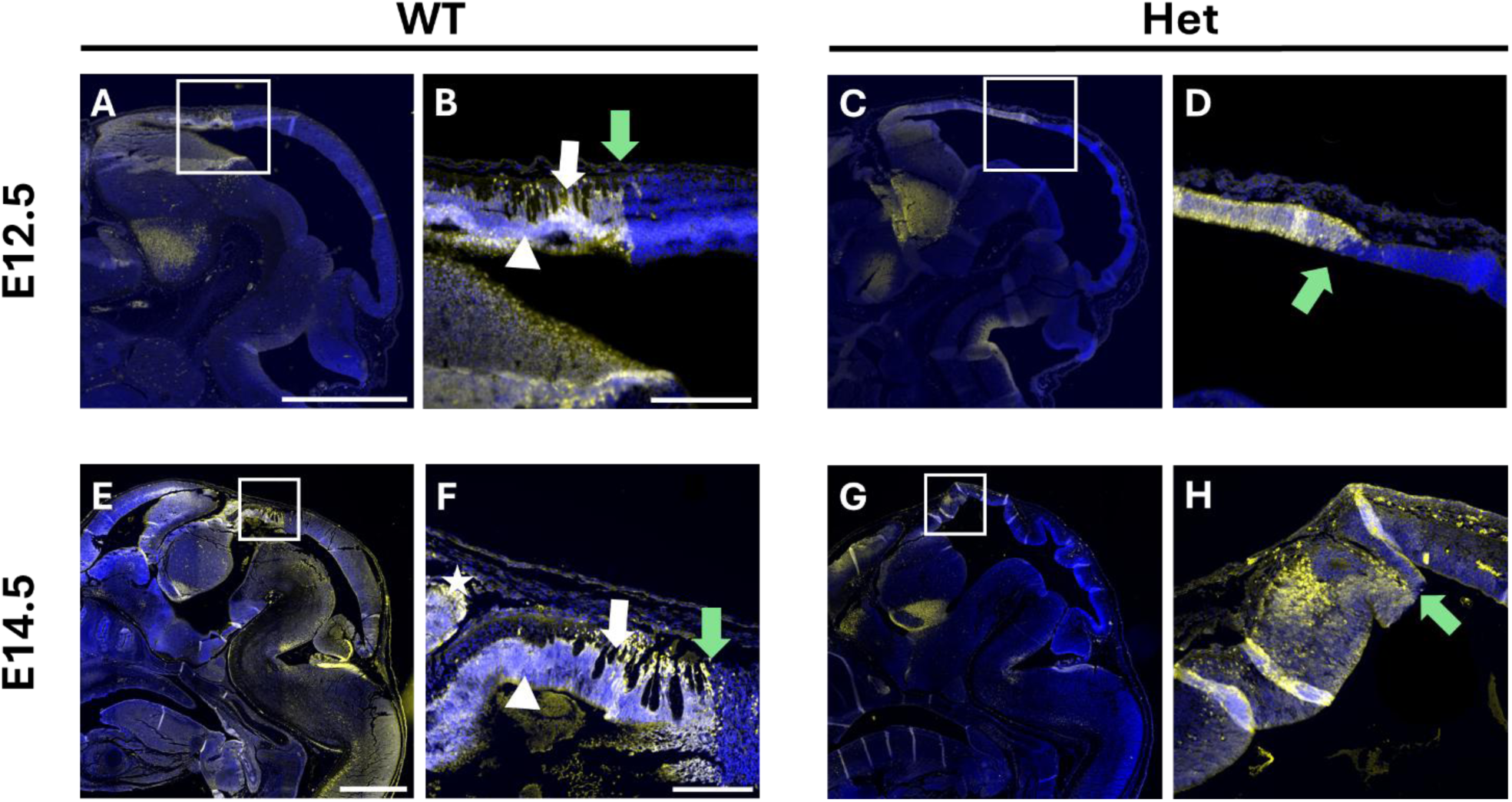
Forebrain-midbrain boundary is correctly specified in pre-encephalocele brains. Immunohistochemistry for Pax6 in mid-sagittal brain sections (anterior to left) of *Wnt1^+/+^; C1-C4^flx/+^* (WT; A,B,E,F) and *Wnt1^Cre/+^; C1-C4^flx/+^* (Het; C,D,G,H) at E12.5 (A-D) and E14.5 (E-H). Boxed areas in A,C,E,G are shown at higher magnification in B,D,F,H. Pax6 expression (white signal over blue DAPI counterstain) is present in the forebrain, as far caudally as the boundary with midbrain (green arrows). WT brains show the posterior commissure (PC; white arrows), sub-commissural organ (SCO; white arrowheads) and pineal gland (asterisk) within the Pax6-positive posterior forebrain region (A,B,E,F). Het embryos show a clear Pax6 expression boundary closely similar to WT embryos, although the PC, SCO and pineal gland are absent at both stages (C,D,G,H). Intense white signal in H is artefactual, due to tissue folds. Scale bars: 0.5 mm (A,C) (E,G); 0.2 mm (B,D) (F,H). Sample size: n = 3 per stage and genotype.

### Change in dimensions of the neuroepithelium and covering skin layer

As well as the absence of a PC-SCO-PG complex from the Het neuroepithelium, we also observed a more general ‘imbalance’ in tissue composition of the dorsal aspect of the embryonic and fetal head. At E12.5 and E14.5, the neuroepithelium of Het embryos and fetuses appears thinner than WT, and frequently exhibits folding internally (Figure 4A-H). Quantification confirmed that, at both E12.5 and E14.5, the Het neuroepithelium is significantly thinner than WT (Figure 4I), has a significantly greater length than WT (Figure 4J), and is significantly reduced in area (length x thickness) (Figure 4K). All three neuroepithelial measures show a significant increase with stage. Hence, the Het neuroepithelium appears ‘attenuated’ compared with WT.

**Figure 4.**
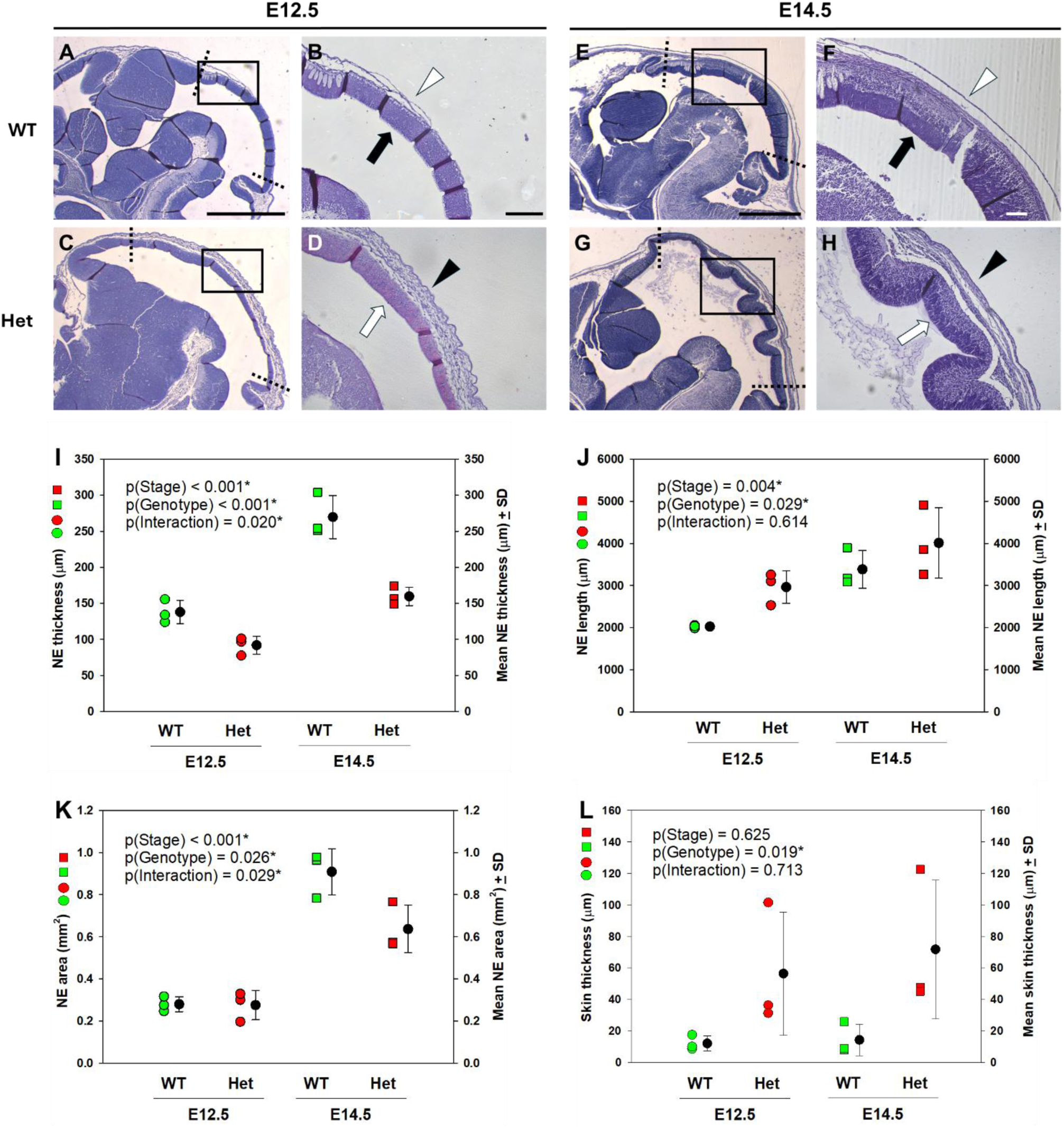
Altered dimensions of neuroepithelium (NE) and covering skin layer in heads of pre-encephalocele embryos and fetuses. **(A-H)** H&E-stained mid-sagittal sections through heads of *Wnt1^+/+^; C1-C4^flx/+^* (WT; A,B,E,F) and *Wnt1^Cre/+^; C1-C4^flx/+^* (Het; C,D,G,H) at E12.5 (A-D) and E14.5 (E-H). Boxed areas in A,C,E,G (or from adjacent sections of the same brains) are shown at higher magnification in B,D,F,H. Note the thinner NE in Het brains at both stages (white arrows in D,H) compared with WT (black arrows in B,F), and the thicker external covering skin layer in Hets (black arrowheads in D,H) compared with WT (white arrowheads in B,F). The Het NE is thrown into folds at E14.5 (G,H). **(I-L)** Quantification of thickness (I), length (J) and sectional area (K) of NE, and thickness of the covering skin layer (L), as measured from mid-sagittal H&E sections. NE length was measured between morphological landmarks indicated by dotted lines in A,C,E,G. Symbols on graphs: WT (green); Het (red); E12.5 (circles); E14.5 (squares). Statistical analysis (2-way ANOVA) shows significant variation (asterisks) by stage and genotype for NE thickness, length and area, whereas covering skin layer thickness varies by genotype but not stage. P-values are shown on the graphs. Scale bars: 0.25 mm (A,C) (B,D); 0.125 mm (E,G) (F,H). Sample size: n = 3 per stage and genotype.

In contrast to these changes in the neuroepithelium, the external skin layer, overlying the posterior forebrain and midbrain, appears markedly thicker in Het than WT (Figure 4A-H). This covering layer comprises the non-neural/surface ectoderm (future epidermis) and, increasingly with developmental stage, mesenchyme cells of neural crest and mesodermal origin, that migrate between the neural and non-neural ectoderm to form dermis, vasculature and bony skull elements. Quantification of skin layer thickness revealed a significant increase in Hets compared with WT. Indeed, the covering layer is 2-10 times thicker in Hets than WT, with considerable variability between Het embryos (Figure 4L). There is no significant change in skin layer thickness with developmental stage.

We conclude that pre-encephalocele embryos, lacking the *C1-C4* insulator sequences upstream of *FGF3, 4 and 15* genes [13], exhibit two types of brain structural defect: a localised absence of the PC-SCO-PG complex and a more generalised abnormality of midline tissue composition, with an abnormally thin neuroepithelium and an excessively thick covering skin layer.

## DISCUSSION

In this study, we investigated the origin of brain defects that culminate in the appearance of parieto-occipital encephalocele in the *Wnt1^Cre/+^; C1-C4^flx/+^* mouse embryo and fetus. The pre-encephalocele phenotype can be identified from E11.5 as a bulging of the head around the midbrain region, and this resolves into a more focal herniation phenotype from E15.5 onwards, in which brain tissue can be seen within the herniated sac. Structural differences are identified between wild-type and mutant embryos and fetuses across different stages, with a main finding of absence from mutant brains of the posterior commissure (PC), sub-commissural organ (SCO) and pineal gland (PG) at all stages studied. Nevertheless, the *Pax6* expression pattern is normal, suggesting that absence of the PC-SCO-PG complex is not due to faulty specification of the forebrain-midbrain boundary. The thickness of the neuroepithelium is reduced in the posterior forebrain and midbrain regions of mutants, whereas the covering skin layer is markedly thickened, suggesting a more global abnormality of head structure that simply an absence of the PC-SCO-PG complex. Hence, a new genetic model of encephalocele demonstrates for the first time fundamental abnormalities in brain midline development, preceding the herniation phenotype.

### Encephalocele as a post-neurulation defect

Encephalocele is usually categorised as an NTD, although the presence of well-formed (‘neurulated’) brain tissue in the herniated sac [18], and the epidemiological characteristics of encephalocele [19] have long suggested a different causation and pathogenesis from the open NTDs: anencephaly and myelomeningocele. Our observations on the *Wnt1^Cre/+^; C1-C4^flx/+^* mouse model confirm that cranial neural tube closure is completed in these mice, prior to the appearance of brain herniation. Previously, we found that mice of genotype *Grhl3^Cre^/Rac1^flox/flox^* develop parieto-occipital encephalocele with rupture of the future epidermal layer that normally covers the closed brain. This leads to brain herniation and a later-arising skull defect overlying the herniated region [20]. However, this mouse model is complex, as embryos with encephalocele co-exist in litters with embryos of identical genotype that exhibit the neural tube closure defect exencephaly. Moreover, most mutant embryos in these litters also develop open spina bifida [20]. This contrasts with the present study in which encephalocele occurs in either *Wnt1^Cre^; C1-C4^flx^* or *Pax3^Cre/+^; C1-C4^flx/+^* embryos as a sole brain phenotype, providing a model that may mimic more closely the human condition of parieto-occipital encephalocele.

### Developmental roles of PC-SCO-PG complex

The PC and SCO are visible in WT brains from E11.5, and the PG is present from E13.5. In contrast, *Wnt1^Cre/+^; C1-C4^flx/+^* and *Pax3^Cre/+^; C1-C4^flx/+^* brains completely lack the entire PC-SCO-PG complex, and show no signs of these structures at any stage. The PC, SCO and PG develop in close proximity to each other, within the posterior forebrain roof plate, and are often jointly affected in mouse mutants, suggesting they have a shared development origin and regulation [21]. The SCO is one of the circumventricular organs that surround the third and fourth ventricles. It consists of ependymal cells that secrete SCO-spondin, later forming Reissner’s fibres, which are suspected of playing an important role in regulation of cerebrospinal fluid flow [22]. Lack or abnormality of the SCO has been associated with brain disorders, particularly hydrocephalus: for example, we found an abnormality of the SCO in E15.5 mouse fetuses with *Gldc* gene loss-of-function, prior to development of ventriculomegaly and hydrocephalus [23]. To our knowledge, however, ours is the first demonstration that lack of PC-SCO-PG can precede appearance of encephalocele.

### Possible mechanisms underlying encephalocele in the mouse model

*Pax6* marks the boundary between forebrain and midbrain [16], and we detected an apparently normal expression pattern of Pax6 in our mutant embryos. This indicates that misexpression of FGF3, 4 and 15 in our encephalocele model does not affect Pax6 expression nor specification of the forebrain-midbrain boundary. While absence of the PC-SCO-PG complex is a striking finding, this is not unique to our model. For example, homozygous *small-eye* mouse mutants, which lack Pax6, also show absence of PC-SCO-PG development [17]. These Pax6-null fetuses exhibit severe brain anomalies and die at birth but, importantly for our study, they do not show an encephalocele phenotype.

These considerations strongly suggest that additional brain abnormalities must be present in *Wnt1^Cre/+^; C1-C4^flx/+^* and *Pax3^Cre/+^; C1-C4^flx/+^* embryos, leading to brain herniation and encephalocele. Indeed, we identified anomalies in the neuroepithelium and covering skin layer of the midbrain dorsal midline that extend beyond the location of the PC-SCO-PG complex. This involves reduced thickness and increased rostro-caudal length of the neuroepithelium, with this layer frequently thrown into folds within the brain ventricle. Moreover, the covering skin layer, which is a simple squamous epithelium in WT embryos, is dramatically thickened in mutants. It remains to be determined whether these tissue changes are direct effects of FGF3, 4, 15 misexpression, or rather are secondary consequences of increased intraventricular pressure in the mutant brain, as suggested by the initial swelling of the midbrain region. The converse is also possible: swelling may occur due to reduced resistance to deformation by the dysmorphic neuroepithelium and covering skin layer.

A further question relates to how the midbrain swelling transitions into a focal brain herniation. This is associated with incomplete formation of the parietal bone [13], which develops from cranial mesoderm via intramembranous bone formation [24]. However, it is unclear whether this bone defect is a direct effect of FGF3, 4, 15 misexpression, or a secondary effect of the brain swelling, which may occupy space needed for formation of the complete parietal bone. The availability of the *Wnt1^Cre/+^; C1-C4^flx/+^* mouse model offers an opportunity to answer these and other questions relating to the pathogenesis of parieto-occipital encephalocele.

## Conclusion

Encephalocele is becoming accepted as a post-neurulation defect of neural tube development, distinct from anencephaly and myelomeningocele. Its presence in mouse embryos and fetuses of the *Wnt1^Cre/+^; C1-C4^flx/+^* and *Pax3^Cre/+^; C1-C4^flx/+^* genotypes provides one of the first experimental models for analysis of this frequently encountered congenital brain disorder. Misexpression of the FGF3, 4 and 15 genes appears to initiate a cascade of developmental events, that leads to midbrain expansion, failure of PC-SCO-PG complex, alterations in the structure of the roof plate neuroepithelium and its covering skin layer, faulty parietal bone development and, finally, herniation of brain tissue within the encephalocele sac. Putting these events into a cause-and-effect pathogenic sequence is an exciting challenge for future research.

## MATERIALS AND METHODS

### Mouse encephalocele model

Mouse research was reviewed and approved by the Animal Welfare and Ethical Review Body of University College London (UCL), and authorised by Project Licence PP0411055 under the auspices of the UK Animals (Scientific Procedures) Act 1986. The mouse encephalocele model was first generated by deletion of a 23.9 kb cluster of four CTCF (CCCTC-binding factor) sites (named *C1-C4*) located upstream of the *Fgf3, 4* and *15* genes on mouse chromosome 7 [13]. The 23.9 kb region was then replaced by a 672 bp region containing only the four CTCF motifs flanked by loxP sites. Mice homozygous for this *C1-C4^flx^* allele were viable and fertile, and offspring in which one copy of the *C1-C4^flx^* allele was removed by *E2A-Cre^t/t^* exhibited the typical encephalocele phenotype [13].

### Mouse procedures

Mice carrying the *C1-C4^flx^* allele were imported into the UCL animal facility and maintained as a homozygous breeding colony. For experiments, *Wnt1^Cre/+^* [25] or *Pax3^Cre/+^* [26] males were mated overnight with homozygous *C1-C4^flx/flx^* females, with the day of finding a copulation plug designated E0.5. Cre-positive offspring (*Wnt1^Cre/+^; C1-C4^flx/+^* or *Pax3^Cre/+^; C1-C4^flx/+^*) exhibited fully penetrant encephalocele, while normally developing (non-Cre; *C1-C4^flx/+^*) littermates served as controls. Genotyping was performed on embryonic or yolk sac DNA, using primers and conditions as previously described for generic *Cre* [27]. During establishment of the breeding colony, *C1-C4^flx^* genotyping used WT forward primer: GCT TTT GAA AGG GAC AGT CAC G; WT reverse primer: CCA ACT AGG TGA CAA GTG GCA GA; floxed allele forward primer: GGA CTC TAG GCG GTG AGA AG; floxed allele reverse primer: CAC AGC TCT CTT GGC TCA GG. These generated a WT band of 155 bp and a floxed allele band of 253 bp. PCR conditions were: 94°C, 2 min, followed by 30 cycles of 94°C, 30 sec; 65°C, 30 sec; 72°C, 30 sec, then 72°C, 2 min, with final hold at 10°C.

### Collection and processing of embryos and fetuses

Embryos and fetuses were dissected from the uterus of pregnant females on each day from E9.5 to E19.5. After removal of extraembryonic membranes, the dissected embryos and fetuses, or isolated heads, were fixed overnight in 10% neutral buffered formalin (CellSolv, #03827005), then washed twice in PBS and dehydrated through an ethanol series: 25%, 50%, 70%, 95%, 100%. After clearing in xylene, whole embryos or embryonic and fetal heads were embedded in paraffin wax (56°C melting point) and sectioned in the sagittal plane on a rotary microtome at 10 µm thickness. Sections on glass slides were dried in an oven overnight before staining.

### Histology

Sections were de-waxed in HistoClear (Geneflow Limited, #A2-0101) followed by hydration through an ethanol series: 100%, 75%, 50%, 25%. Sections were stained with Harris Haematoxylin solution (Sigma Aldrich, #0000259604) and Eosin Y solution (Sigma Aldrich, #0000288375). Sections were quickly dehydrated through the ethanol series, immersed in HistoClear, and then mounted in DPX (Thermo Fisher Scientific, #1731605). After overnight drying, images were taken on a Zeiss Axioplan microscope.

### Immunofluorescence

Tissue sections were de-waxed in HistoClear, rehydrated through an ethanol series: 100%, 70%, 50%, 25%, and washed under running tap water. The sections were then fully immersed in antigen retrieval solution [sodium citrate tribasic dihydrate (Sigma-Aldrich, #BCCC4347), Tween® 20 (Sigma-Aldrich, #SLCC6910), 1X PBS], and placed in a steamer for 10 min, followed by 0.1% PBS-T washes [1X PBS, Triton X-100 (Sigma-Aldrich, #T8787)]. Sections were blocked for 1 h in blocking solution [5% bovine serum albumin (Sigma-Aldrich, #SLCD3068), 0.1% PBS-T]. The sections were incubated with rabbit anti-Pax6 primary antibody (BioLegend, #901301), diluted 1:100 in blocking solution, at 4°C overnight in a dark, humidified chamber. Sections were washed in 0.1% PBS-T to remove excess primary antibody, and then incubated with Alexa Fluor 488 goat anti-rabbit secondary antibody (Invitrogen, #A11008), diluted 1:200 in blocking solution, at room temperature in a dark, humidified chamber. DAPI staining (1:10000, Severn Biotech Ltd, #30-45-01) for 10 min was followed by 0.1% PBS-T washes. Slides were mounted using ProLong Gold antifade reagent (Invitrogen, #2696990), sealed with nail polish and cured overnight in the dark at room temperature. Images were captured on a Zeiss Observer microscope.

### Image analysis and statistics

Neuroepithelial and covering (skin) layer thickness were measured using ImageJ (protocol available upon request). Neuroepithelial length measurements were made between anatomical landmarks, as indicated by the dashed lines in Figure 4A,C,E,G. Neuroepithelial area was calculated as thickness x length. For neuroepithelium and covering skin layer measurements, 8 sections were quantified for each embryo, with mean values used as biological replicates. For immunofluorescence staining, 4 sections were analysed per embryo. Neuroepithelium and covering skin layer dimensions were compared between WT and Het, at E12.5 and E14.5, by two-way analysis of variance (genotypes, stages), computed using Sigmaplot v16.

## Acknowledgements

The authors thank Smiles with Grace and Great Ormond Street Hospital Charity (grant number W1172) for financial support.

